# SARS-CoV-2 launches a unique transcriptional signature from in vitro, ex vivo, and in vivo systems

**DOI:** 10.1101/2020.03.24.004655

**Authors:** Daniel Blanco-Melo, Benjamin E. Nilsson-Payant, Wen-Chun Liu, Rasmus Møller, Maryline Panis, David Sachs, Randy A. Albrecht, Benjamin R. tenOever

**Affiliations:** Department of Microbiology, Icahn School of Medicine at Mount Sinai, New York, USA; Virus Engineering Center for Therapeutics and Research (VECToR), Icahn School of Medicine at Mount Sinai, New York, USA; Global Health and Emerging Pathogens Institute, Icahn School of Medicine at Mount Sinai, New York, USA; Department of Genetics and Genomic Sciences, Icahn School of Medicine at Mount Sinai, New York, USA

## Abstract

One of the greatest threats to humanity is the emergence of a pandemic virus. Among those with the greatest potential for such an event include influenza viruses and coronaviruses. In the last century alone, we have observed four major influenza A virus pandemics as well as the emergence of three highly pathogenic coronaviruses including SARS-CoV-2, the causative agent of the ongoing COVID-19 pandemic. As no effective antiviral treatments or vaccines are presently available against SARS-CoV-2, it is important to understand the host response to this virus as this may guide the efforts in development towards novel therapeutics. Here, we offer the first in-depth characterization of the host transcriptional response to SARS-CoV-2 and other respiratory infections through *in vitro, ex vivo*, and *in vivo* model systems. Our data demonstrate the each virus elicits both core antiviral components as well as unique transcriptional footprints. Compared to the response to influenza A virus and respiratory syncytial virus, SARS-CoV-2 elicits a muted response that lacks robust induction of a subset of cytokines including the Type I and Type III interferons as well as a numerous chemokines. Taken together, these data suggest that the unique transcriptional signature of this virus may be responsible for the development of COVID-19.

## INTRODUCTION

Coronaviruses are a diverse group of single-stranded positive-sense RNA viruses infecting a wide range of vertebrate hosts^1^. These viruses are thought to generally cause mild upper respiratory tract illnesses in humans such as the common cold^2^. However, in the past two decades, three highly pathogenic human coronaviruses have emerged from zoonotic viruses: severe acute respiratory syndrome-related coronavirus (SARS-CoV-1) infecting ∼8,000 people worldwide with a case-fatality rate of ∼10% in 2002-2003, Middle East respiratory syndrome-related coronavirus (MERS-CoV) infecting ∼2,500 people with a case-fatality rate of ∼36%, and now severe acute respiratory syndrome-related coronavirus 2 (SARS-CoV-2) which causes Coronavirus Disease-2019 (COVID-19) whose global mortality rate remains to be determined^3, 4^. Infection with these highly pathogenic coronaviruses can result in acute respiratory distress syndrome (ARDS) and acute lung injury (ALI), often leading to reduction of lung function and even death^3^.

The current pandemic of COVID-19 represents an acute and rapidly developing global health crisis. In an effort to better understand the molecular basis of the disease and identify putative markers for COVID-19, we compared the transcriptional response of SARS-CoV-2 to that of seasonal influenza A virus (IAV) and human orthopneumovirus (commonly known as human respiratory syncytial virus (RSV)), two common recurring respiratory viruses. Comparing the transcriptional response in both primary human lung epithelium and transformed lung alveolar cells revealed that SARS-CoV-2 elicits a unique transcriptional response as compared to IAV and RSV. In a homogenous cell population, the transcriptional response to SARS-CoV-2 infection shows a significant lack of Type I and III interferon (IFN-I and IFN-III) expression as compared to IAV and RSV. Moreover, while a core number of cytokines that comprise the antiviral host defense are shared amongst all three viruses, an equal number are notably absent in response to SARS-CoV-2. Lastly, the only genes that appear to be unique to SARS-CoV-2 infection are secreted peptides implicated in diseases of the airways. This unique response to SARS-CoV-2 could also be recapitulated *in vivo* comparing influenza infection to COVID-19 in ferrets. Taken together, these data reveal that the interaction between virus and host as it relates to SARS-CoV-2 may be responsible for its unusual high morbidity and mortality.

## RESULTS

In order to identify similarities and differences between the host response to SARS-CoV-2 and seasonal respiratory infections, we sought to elucidate the transcriptome of lung epithelial cells during viral infection. To achieve this, we infected human alveolar adenocarcinoma cells with SARS-CoV-2, RSV and IAV and performed mRNA-seq analysis (Supplementary Table 1). Despite undetectable levels of ACE2 and TMPRSS2, the putative receptor and protease for SARS-CoV-2 entry^5^, these cells were able to support viral replication, evidenced by full genome coverage in total RNA samples extracted from infected cells (Figure 1a and Extended Data Fig.1a). Our analysis identified 120 differentially expressed genes (DEGs, qval < 0.05) of which more than 80% were up-regulated (Figure 1b and Supplementary Table 2). Analysis of DEGs revealed two main clusters of induced genes (Figure 1c). A highly interacting cluster enriched in genes involved in the cellular response to virus infection (GO:0009615, FDR << 0.0001), mainly composed of type-I ISGs (GO:0034340, FDR << 0.0001), and a second cluster enriched in genes involved in the humoral immune response (GO:0006959, FDR << 0.0001), which further subdivided into two smaller clusters enriched in chemokines and cytokines (GO:0005125, FDR < 0.001), and complement proteins (GO:0006956, FDR < 0.0005) (Figure 1c and Supplementary Table 3). It is noteworthy to highlight that, despite a lack of IFN-I and IFN-III expression, we observe the induction of well-characterized direct effectors of the innate immune response including: MX1, IFITM3, SAMHD1 and TRIM25, as well as the induction of viral RNA sensors such as RIG-I and the OAS1-3 genes (Figure 1c). This response however is lacking the robust induction of antiviral genes commonly observed following IFN-I/-III signaling^6^. This muted response however does show a positive correlation in the host antiviral response overall as determined by gene set enrichment analysis (GSEA) (Extended Data Fig. 1b). Taken together our analyses indicate that replication of SARS-CoV-2 in alveolar cells results in a limited antiviral response.

**Figure 1.**
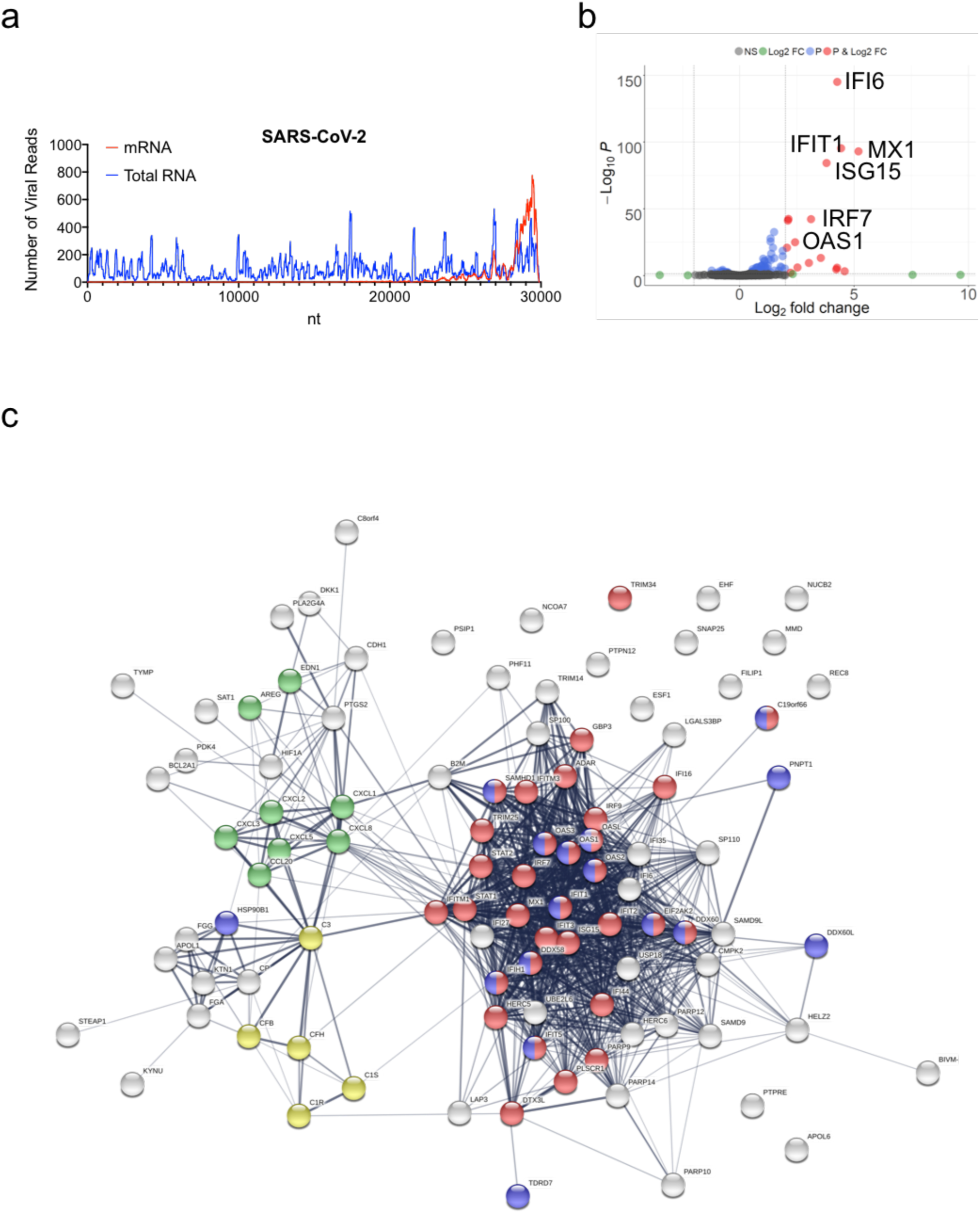
Host Transcriptional response to SARS-CoV-2 infection in A549 cells. (**a**) read coverage along the SARS-CoV-2 genome. Number of viral reads per each position of the virus genome. Blue graph indicate read coverage when NGS libraries were prepared using Ilumina’s TruSeq Stranded Total RNA Gold kit. Red graph indicate read coverage when NGS libraries were prepared using Ilumina’s TruSeq RNA Library Prep Kit v2. (**b**) Volcano plot depicting differentially expressed genes in response to SARS-CoV-2 infection. Red dots indicate genes with a |Log_2_(Fold Change)| > 2. The identity of top induced genes is indicated. (**c**) Protein interaction network of significantly induced genes in response to SARS-CoV-2 infection. Genes involved in enriched biological processes and molecular functions are indicated in color. Red: genes involved in the response to virus (GO:009615, FDR<<0.0001). Green: genes with cytokine activity (GO:0005125, FDR<0.001). Yellow: genes involved in complement activation (GO:0006956, FDR<0.0005). Blue: genes with RNA binding capability (GO: 0003723, FDR<0.0005). Gene enrichment analyses were performed using STRING.

In an effort to determine if the apparent modest response to SARS-CoV-2 infection was the result of low receptor expression, a low multiplicity of infection (MOI), or due to the immortalized nature of the cell line, we next infected primary human bronchial epithelial (NHBE) cells (Fig. 2a). However, despite a ten fold increase in MOI, infection of NHBE cells showed a similar transcriptional response to that of the immortalized alveolar cells (ρ≈0.22, p-value << 0.0001). This correlation extended to both the type-I IFN response (GO:0060337, FDR << 0.0001) as well as the humoral immunity (GO:0006959, FDR << 0.0001) (Fig. 2b). Taken together, these results suggest that the cellular response to SARS-CoV-2 is relatively uniform and void of IFN-I and IFN-III expression.

**Figure 2:**
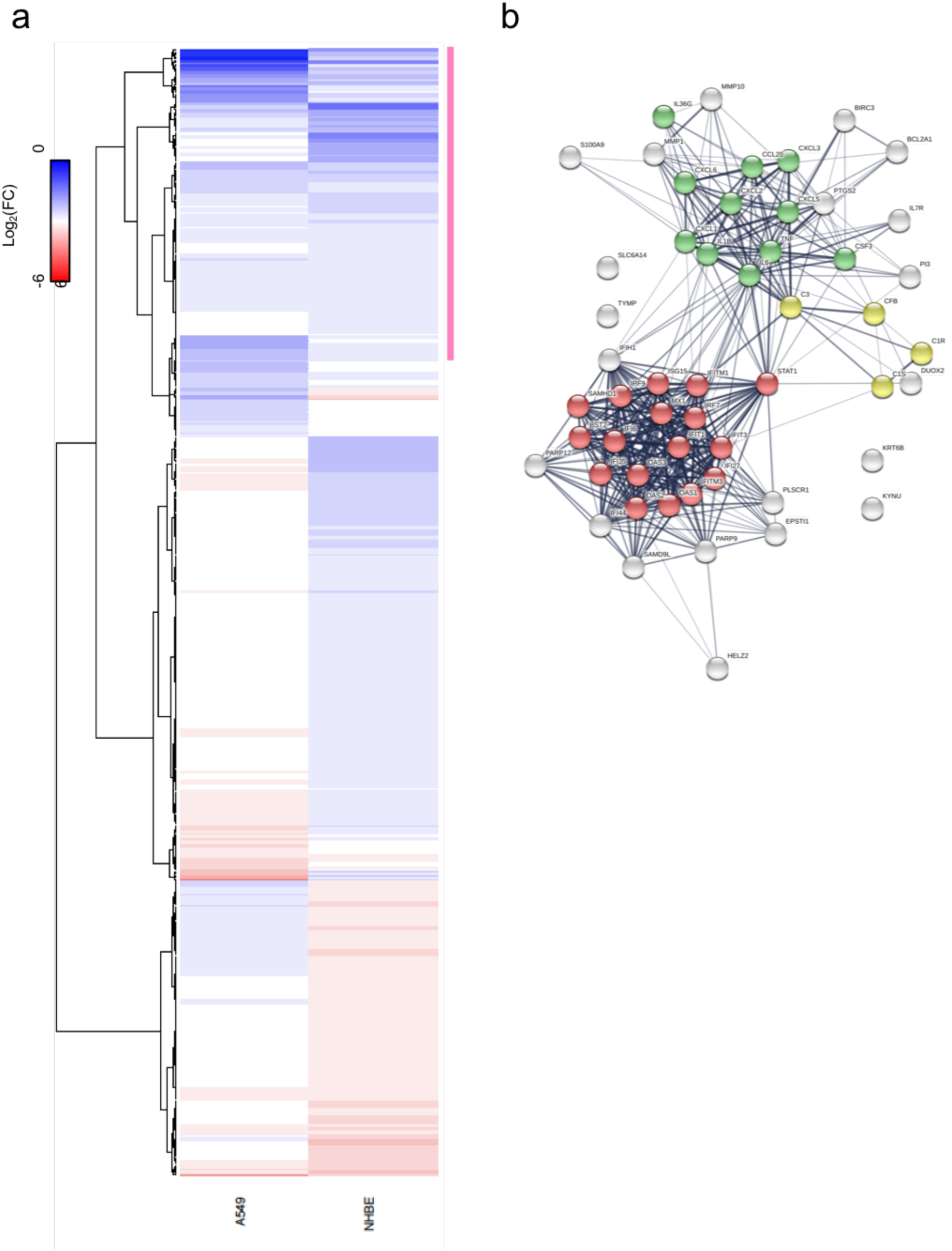
Host Transcriptional response to SARS-CoV-2 infection in NHBE cells. (**a**) Expression levels of DEGs in response to SARS-CoV-2 infection in A549 and NHBE cells. Heatmap depicting the expression of DEGs in both cell types. Pink bar indicates a cluster of similarly induced genes upon SARS-CoV-2 infection in both cell types. (**b**) Protein interaction network of genes indicated in pink in (C) in response to SARS-CoV-2 infection. Genes involved in enriched biological processes and molecular functions are indicated in color. Red: genes involved in the type-I IFN signaling pathway (GO:0060337, FDR<<0.0001). Green: genes with cytokine activity (GO:0005125, FDR<<0.001). Yellow: genes involved in complement activation (GO:0006956, FDR<0.0005). Gene enrichment analyses were performed using STRING.

In an effort to compare the response to SARS-CoV-2 with other respiratory viruses, we next infected the lung alveolar carcinoma cell line with either RSV or seasonal IAV. Bulk RNA sequencing of independent biological replicates revealed that the transcriptional response to SARS-CoV-2 infection is similar in magnitude to that of IAV (82 DEGs, qval < 0.05), with both viruses showing a moderate response compared to that of RSV (910 DEGs, qval < 0.05) (Supplementary Table 4 and 5). Nevertheless, the response to SARS-CoV-2 showed a strong correlation (p-value < 0.001) to that of RSV infection (Extended Data Fig. 2), resulting in the large overlap of DEGs between these two viruses, compared to IAV (Fig. 3a). These responses do not reflect the replication dynamics of these viruses, illustrated by the amount of viral reads recovered from these infections (Fig. 3b). In an effort to better illustrate the unique responses to SARS-CoV-2, IAV, and RSV, we next performed a principle component analysis (PCA) to enable unbiased grouping of each sample based in DEGs. These efforts not only highlighted the unique transcriptional signatures induced by the three respiratory infections, they also corroborated that the response in different cell systems with SARS-CoV-2 were comparable (Fig. 3c). To further delineate the unique antiviral signatures of each virus, we next examined the expression levels of specific genes implicated in the host antiviral response (Fig. 3d). These analyses again showed that despite sharing a similar transcriptional footprint to RSV, SARS-CoV-2 lacks the induction of IFN-I and IFN-III genes (Fig. 3d). However, this same analyses shows a common subset of well established interferon stimulated genes (ISGs) shared with RSV that are lacking in IAV, presumably due to the expression of the antiviral antagonist NS1^7^. While most of the transcriptional response to SARS-CoV-2 was either shared or lacking in comparison to the response to IAV and RSV, two cytokines (EDN1 and TNFSF15) were uniquely upregulated (Fig. 3e). The unique induction of these two secreted factors was particularly noteworthy as they have been implicated as mediators of respiratory inflammation^6^.

**Figure 3:**
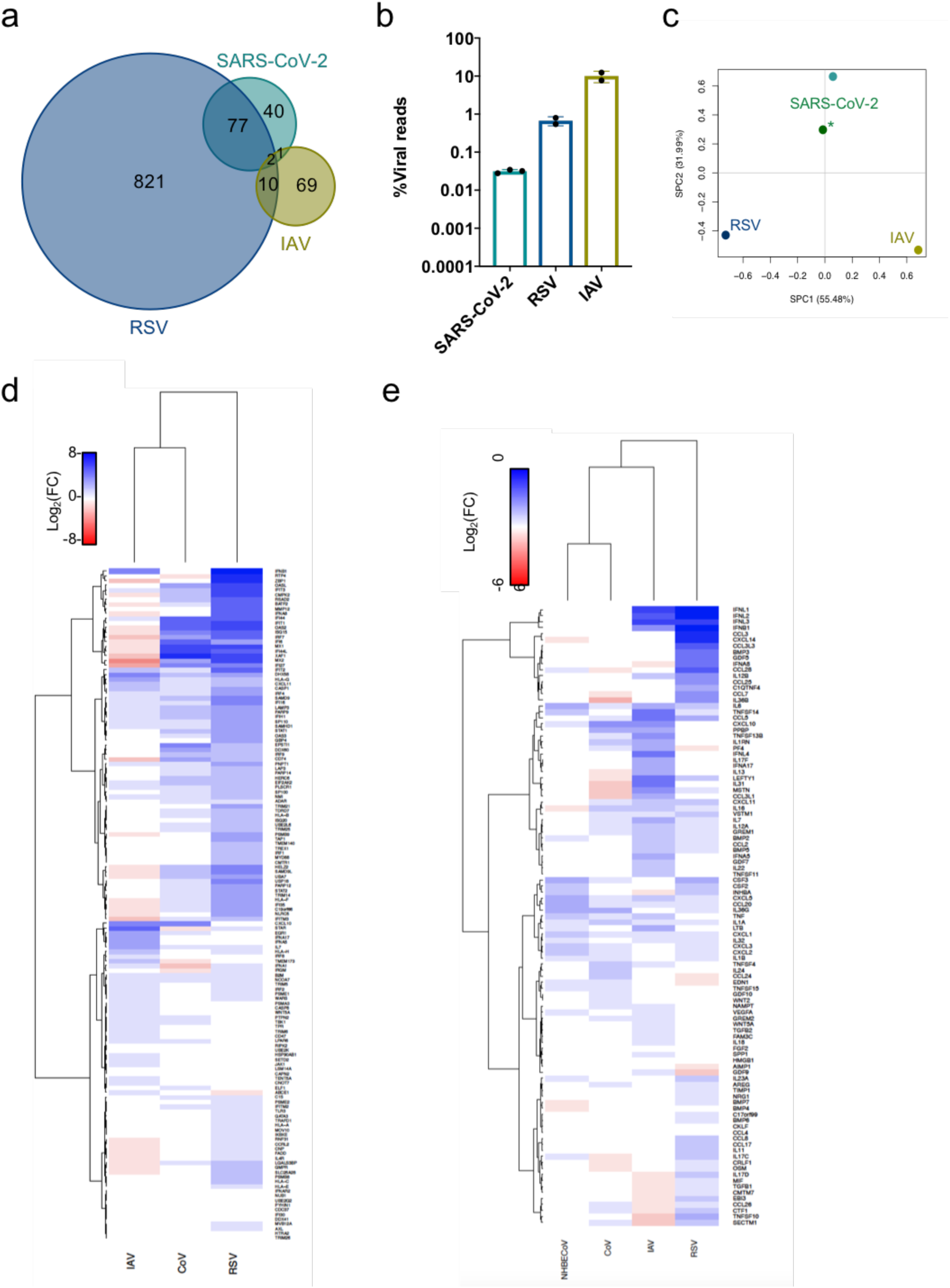
Unique host transcriptional profiles elicited differentially in response to diverse respiratory viruses. (**a**) Shared DEGs in SARS-CoV-2, RSV and IAV infected A549 cells. Venn diagram of DEGs in each sample, genes shared/unique between each sample are indicated. (**b**) Virus replication levels in A549 cells. Percentage of virus-aligned reads (over total reads) for each infected sample. Data from 3 independent biological replicates. (**c**) Principal component analysis (PCA) for the global transcriptional response to respiratory viruses. Sparse PCA depicting global transcriptome profiles between samples. All data from infections in A549 cells, except for (*) that was performed in NHBE cells. (**d-e**) Heatmaps depicting the expression levels of genes involved in the type-I IFN response (d) or with cytokine or chemokine activity (e).

Lastly, to ascertain whether the mutated antiviral response to SARS-CoV-2 could be observed *in vivo*, we infected ferrets with either influenza A virus or SARS-CoV-2 (Fig. 4). At five days post infection, nasal washes were performed and the heterogeneous cells obtained from this procedure were again subjected to bulk RNA sequencing and aligned to either SARS-CoV-2 or A/California/04/2009 (Fig. 4a). These data demonstrate complete coverage of the challenge virus suggesting that in both examples, the ferrets were shedding replication competent virus. Next, these same upper respiratory cell populations were compared to mock treated ferrets to ascertain the transcriptional response *in vivo* for comparison purposes. In agreement with our *in vitro* data, differential expression analyses of these respiratory infections demonstrated that the transcriptional footprint of IAV was significantly greater in magnitude that that observed for SARS-CoV-2 (Fig. 4b-c). While the ferret genome annotation limits the direct comparisons between some orthologues, we do observe a diminished cytokine response when comparing SARS-CoV-2 to IAV. These data include a subset of genes involved in the recruitment and activation of the adaptive immune response including: IL6, CXCL11, IFNG, IL7, CXCL9, and CXCL10 (Fig. 4b-c). Taken together with both our *in vitro* and *ex vivo* data, which also implicated a diminished cytokine response to infection, these results suggest that the overall response to SARS-CoV-2 is relatively moderate when compared to other respiratory viruses.

**Figure 4:**
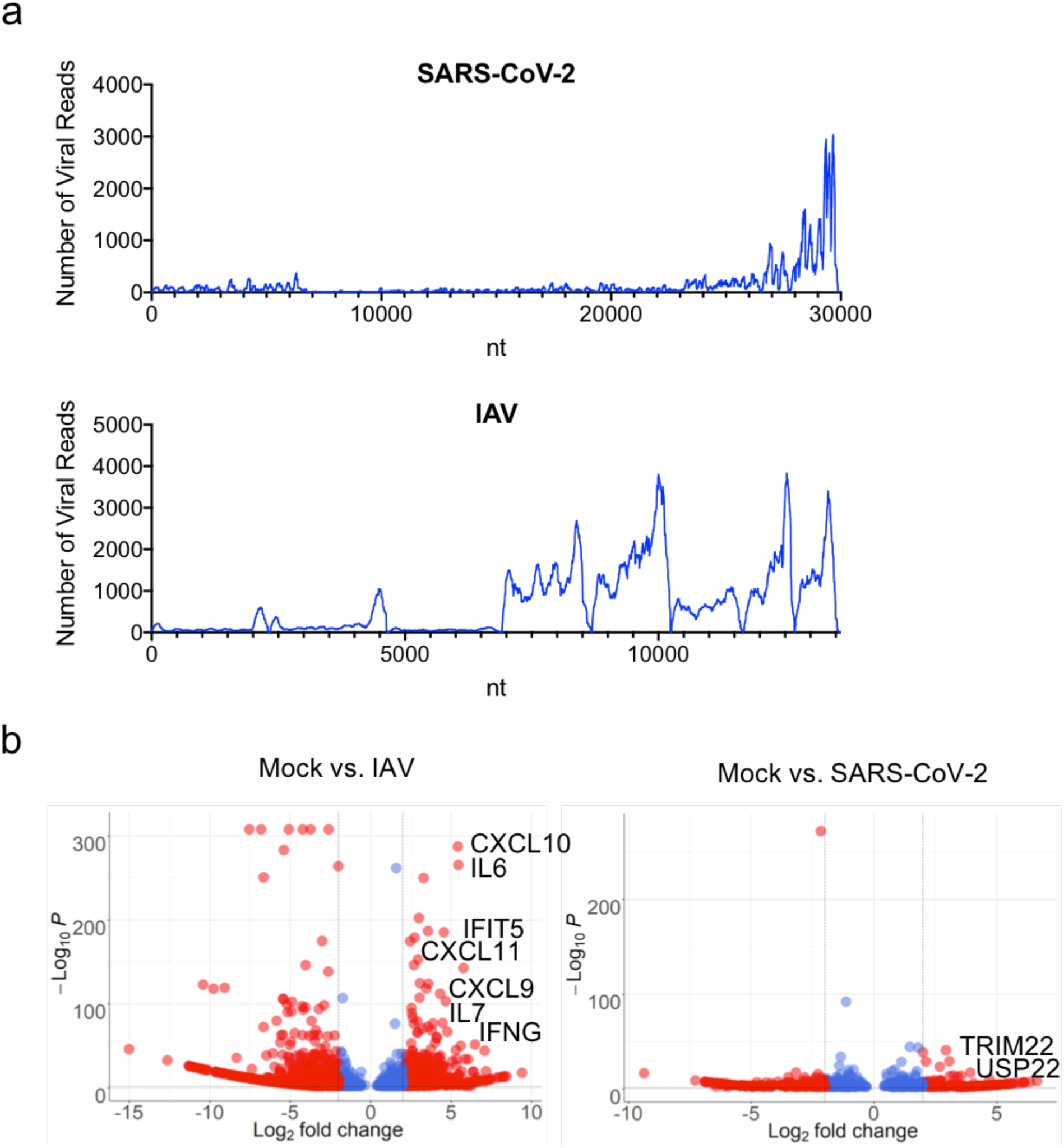
Host Transcriptional response to SARS-CoV-2 infection *in vivo*. (**a**) read coverage along the SARS-CoV-2 and IAV (A/California/04) genomes. Number of viral reads per each position of the virus genome. (**b**) Volcano plot depicting differentially expressed genes in response to SARS-CoV-2 or IAV (A/California/04) infection. Red dots indicate genes with a |Log_2_(Fold Change)| > 2. The identity of top induced genes is indicated.

## DISCUSSION

In the present study we focus on defining the transcriptional response to SARS-CoV-2, influenza A virus, and respiratory syncytial virus. In general, these data find that the overall transcriptional footprint to infection is greatest for RSV and the lowest for IAV with SARS-CoV-2 representing an intermediate response. A probable explanation for the lack of a response to IAV in general is the potent antagonistic activity of the virus. However, it is noteworthy that *in vivo*, the response to IAV exceeds that of SARS-CoV-2 where the activity of NS1 is unable to inhibit some of the pattern recognition receptors at play during a physiological response.

Despite the apparent muted induction of antiviral genes in response to SARS-CoV-2, we do observe a significant up regulation of well-characterized ISGs including: IFIT1-3, ISG15, DDX58, and others. Amongst the genes that are uniquely present when comparing SARS-CoV-2 to other respiratory viruses are EDN1 and TNFSF15 - two putative biomarkers that may contribute to COVID-19 pathology. However, it should be noted that we did not observe these genes in ferrets despite the fact that they could be detected in both cell culture models. Regardless of whether these two unique genes may serve as biomarkers, the overall signature of SARS-CoV-2, RSV, and IAV can serve as a broader map to developing novel diagnostic strategies.

The general induction of ISGs in response to SARS-CoV-2, albeit modest, is particularly interesting as we were not able to detect any reads mapping to IFN-I of IFN-III members in contrast to what is observed for RSV or IAV. These data likely indicate that induction of IFN-I and IFN-III is very low but sufficient to induce at least a subset of ISGs in response to SARS-CoV-2. It is also noteworthy that we observe comparable replication between primary human bronchial epithelium and a transformed cell line despite the fact that we are unable to detect both the putative receptor (ACE2) or the required protease (TMPRSS2) in the latter cell line. While further work remains to ascertain how viral entry is mediated in this model system, we did note high expression of BSG which has also been suggested to compensate for ACE2 (Wang et al, BioRxv unpublished and Extended Data Fig. 1a). Moreover, it should be noted that the viral replication profiles as deduced by both total RNA and polyA RNA sequencing were comparable between *in vitro, ex vivo*, and *in vivo* samples.

What makes the SARS-CoV-2 distinct from the RSV and IAV strains used in this study is the propensity to selectively induce morbidity and mortality in older populations^8^. The physiological basis for this morbidity is believed to be the selective death of Type II pneumocytes that results in both loss of air exchange and fluid leakage into the lungs^9, 10^. While it remains to be determined whether this moderate cell response to SARS-CoV-2 is responsible for the abnormally high lethality in the older populations, it does explain why the virus is generally asymptomatic in young people with healthy and robust immune systems^11^. Given the results here, it is tempting to speculate that perhaps in the aging population, the immune response itself is muted and thus prevents successful inhibition of viral spread. Perhaps the slow amplification of the virus is then the underlying cause for the lung damage and an explanation for why the course of infection is so prolonged. In this regard, it is also of interest to point out that while replication is apparent in 6 month old ferrets (comparable in age to human teenagers), they do not show overt signs of disease and clear infection after approximately eight days. Taken together, the data presented here suggests that perhaps artificial means of boosting the antiviral response may be an effective option at treating COVID-19 so long as it does not further aggravate any pre-existing conditions.

## MATERIALS AND METHODS

### Cell Culture

Human adenocarcinomic alveolar basal epithelial (A549) cells (ATCC, CCL-185), Madin-Darby Canine Kidney (MDCK) cells (ATCC, CCL-34), African green monkey kidney epithelical Vero E6 cells (ATCC, CRL-1586) and African green monkey kidney epithelical BS-C-1 cells (ATCC, CCL-26) were maintained at 37°C and 5% CO_2_ in Dulbecco’s Modified Eagle Medium (DMEM, Gibco) supplemented with 10% Fetal Bovine Serum (FBS, Corning). Normal human bronchial epithelial (NHBE) cells (Lonza, CC-2540 Lot# 580580) were isolated from a 79-year-old Caucasian female and were maintained in bronchial epithelial growth media (Lonza, CC-3171) supplemented with BEGM SingleQuots as per the manufacturer’s instructions (Lonza, CC-4175) at 37°C and 5% CO_2_.

### Viruses

Influenza A/Puerto Rico/8/1934 (H1N1) virus (NCBI:txid183764) and influenza A/California/04/2009 (pH1N1) virus was grown in MDCK cells at an MOI of 0.001 in DMEM supplemented with 0.3% bovine serum albumin (BSA, MP Biomedicals), 4.5 g/L D-glucose, 4 mM L-glutamine and 1 µg/ml TPCK-trypsin (Sigma-Aldrich). Infectious titers of influenza A viruses were determined by plaque assay in MDCK cells.

Recombinant GFP-expressing human respiratory syncytial virus, strain A2 (rgRSV[224]) was generously provided by Dr. M. Peeples (OSU) and was described previously^12^. rgRSV[224] was grown in BSC-1 cells in in DMEM supplemented with 2% FBS, 4.5 g/L D-glucose and 4 mM L-glutamine.

SARS-related coronavirus 2 (SARS-CoV-2), Isolate USA-WA1/2020 (NR-52281) was deposited by the Center for Disease Control and Prevention and obtained through BEI Resources, NIAID, NIH. SARS-CoV-2 was propagated in Vero E6 cells in DMEM supplemented with 2% FBS, 4.5 g/L D-glucose, 4 mM L-glutamine, 10 mM Non-Essential Amino Acids, 1 mM Sodium Pyruvate and 10 mM HEPES. Infectious titers of SARS-CoV-2 were determined by plaque assay in Vero E6 cells in Minimum Essential Media supplemented with 4 mM L-glutamine, 0.2% BSA, 10 mM HEPES and 0.12% NaHCO_3_ and 0.7% agar.

All work involving live SARS-CoV-2 was performed in the CDC/USDA-approved BSL-3 facility of the Global Health and Emerging Pathogens Institute at the Icahn School of Medicine at Mount Sinai in accordance with institutional biosafety requirements.

### RNA-Seq of viral infections

Approximately 1 × 10^6^ A549 cells were infected with influenza A/Puerto Rico/8/1934 (H1N1) virus (IAV), human respiratory syncytial virus (RSV) or SARS-CoV-2. Infections with IAV were performed at a multiplicity of infection of 5 for 9 h in DMEM supplemented with 0.3% BSA, 4.5 g/L D-glucose, 4 mM L-glutamine and 1 µg/ml TPCK-trypsin. Infections with RSV were performed at an MOI of 15 for 24 h in DMEM supplemented with 0.3% BSA, 4.5 g/L D-glucose and 4 mM L-glutamine. Infections with SARS-CoV-2 were performed at an MOI of 0.2 for 24 h in DMEM supplemented with 2% FBS, 4.5 g/L D-glucose, 4 mM L-glutamine, 10 mM Non-Essential Amino Acids, 1 mM Sodium Pyruvate and 10 mM HEPES. Approximately 1 × 10^5^ NHBE cells were infected with SARS-CoV-2 at an MOI of 2 for 24 h in bronchial epithelial growth media supplemented with BEGM SingleQuots.

Total RNA from infected and mock infected cells was extracted using TRIzol Reagent (Invitrogen) and Direct-zol RNA Miniprep kit (Zymo Research) according to the manufacturer’s instructions and treated with DNase I.

RNA-seq libraries of polyadenylated RNA were prepared using the TruSeq RNA Library Prep Kit v2 (Illumina) according to the manufacturer’s instructions. RNA-seq libraries for total ribosomal RNA-depleted RNA were prepared using the TruSeq Stranded Total RNA Library Prep Gold (Illumina) according to the manufacturer’s instructions. cDNA libraries were sequenced using an Illumina NextSeq 500 platform.

### Bioinformatic analyses

Raw reads were aligned to the human genome (hg19) using the RNA-Seq Aligment App on Basespace (Illumina, CA), following differential expression analysis using DESeq2^13^. Differentially expressed genes (DEGs) were characterized for each sample (p adjusted-value < 0.05) and were used as query to search for enriched biological processes (Gene ontology BP) and network analysis of protein interactions using STRING^14^. Heatmaps of gene expression levels were constructing using heatmap.2 from the gplot package in R (https://cran.r-project.org/web/packages/gplots/index.html). Sparse principal component analysis (sPCA) was performed on Log_2_(Fold Change) values using SPC from the PMA package in R^15^. Volcano plots were constructed using custom scripts in R. Gene set enrichment analysis (GSEA) was performed on gene expression profiles from mock and SARS-CoV-2 treated cells comparing to a composite list of genes involved in Type-I IFN response (GO:0035457, GO:0035458, GO:0035455, GO:0035456, GO:0034340, and HALLMARK_INTERFERON_ALPHA_RESPONSE)^16^. Correlation analysis on DEGs between samples was performed using chart.Correlation from the PerformanceAnalytics package in R (https://cran.r-project.org/web/packages/PerformanceAnalytics/index.html). Alignments to viral genomes was performed using bowtie2. The genomes used for this study were: SARS-CoV-2 (NC_045512.2), RSV (NC_001803.1), IAV PR8 (AF389115.1, AF389116.1, AF389117.1, AF389118.1, AF389119.1, AF389120.1, AF389121.1, AF389122.1) and IAV A/California/VRDL6/2010(H1N1) (CY064994, CY064993, CY064992, CY064987, CY064990, CY064989, CY064988, CY064991).

#### Ferret immunization and infection

All procedures are described in our previous study^17^. Briefly, outbred 4-month old castrated male Fitch ferrets were confirmed to be seronegative for circulating H1N1, H3N2, and B influenza viruses prior to purchase from Triple F. Farm (North Rose, NY). All animal experiments were performed according to protocols approved by the Institutional Animal Care and Use Committee (IACUC) and Institutional Biosafety Committee of the ISMMS. Ferrets were randomly assigned to the different treatment groups. For each experiment, the experimental group size is indicated in the figure legends. For pH1N1 virus infection, all naive ferrets were intranasally challenge infected with 1mL of 10^5^ PFU of pH1N1 (A/California/04/2009) influenza virus. Nasal wash samples were collected from anesthetized ferrets on day 7 post infection and preserved at −80°C for viral titration. Cell lysates were treated with Trizol for bulk RNA sequencing. For the SARS-CoV-2 virus infection study, all naïve ferrets were infected with 1mL of 5×10^4^ PFU of the SARS-CoV-2 isolate USA-WA1/2020. Nasal wash samples were collected from anesthetized ferrets on day 5 post-infection, and preserved at −80 °C for viral titration. Cell lysates were treated with Trizol for bulk RNA sequencing. On day 5 post-challenge infection, anesthetized ferrets were euthanized by exsanguination followed by intracardiac injection of euthanasia solution (Sodium Pentobarbital). Trachea samples were collected from each ferret (n=3) to quantify viral titers by plaque assays and for bulk RNA sequencing.

## Acknowledgments

DBM is an Open Philanthropy Fellow for the Life Sciences Research Foundation. BT and this work are supported by the Defense Advanced Research Projects Agency (DARPA). For raw RNA seq. data, please contact

---

**Extended Data Figure 1:**
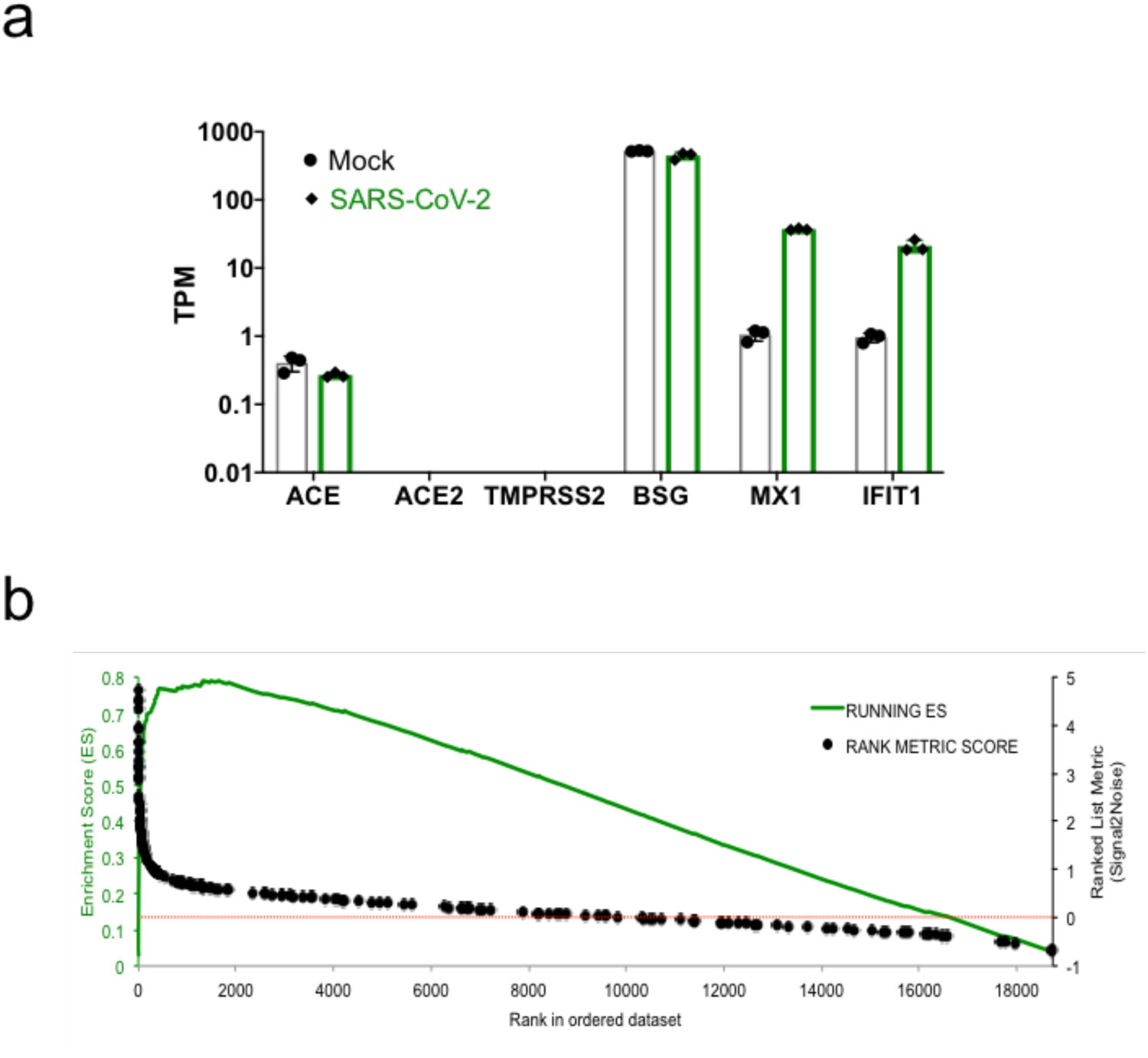
Host Transcriptional response to SARS-CoV-2 infection (Related to figure 1). (**A**) Expression levels of distinct genes in mock treated or SARS-CoV-2 infected A549 cells. TPM: transcript per million. Data from 3 independent biological replicates. (**B**) Gene set enrichment analysis (GSEA) of genes involved in the type-I IFN response. Gene Set was constructed of a composite of type-I IFN gene sets (see methods). Ranked genes based of the gene expression levels of SARS-CoV-2 infected A549s compared to Mock treated cells are depicted in the horizontal axis. Green line depicts the running enrichment score along the ranked genes. Black dots represent the location (and metric score) of type-I IFN related genes along the ranked genes. A doted red line represents the transition between SARS-CoV-2 (>0) and mock (<0) correlated genes.

**Extended Data Figure 2:**
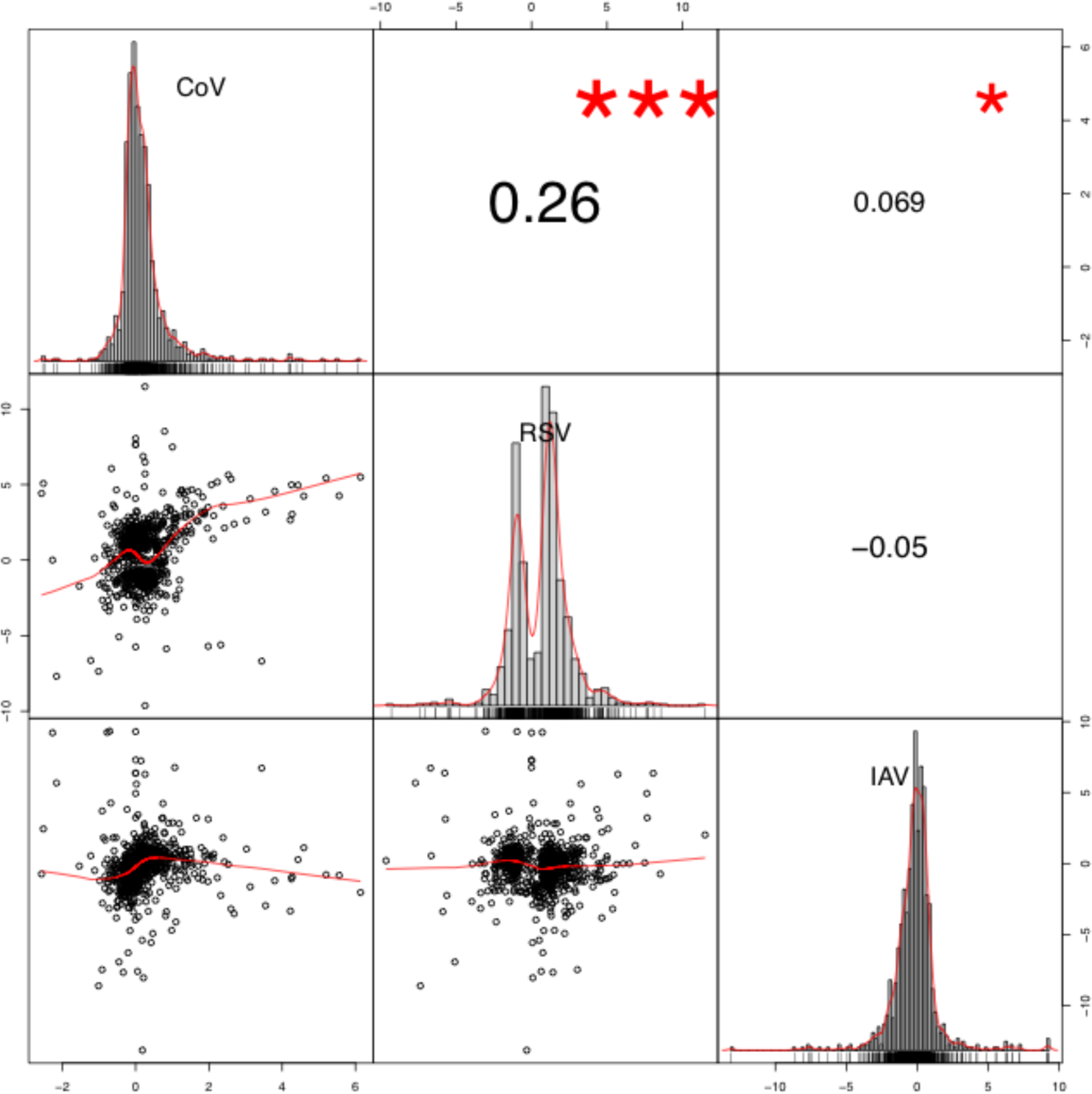
Comparison of the host response to respiratory viral infections. (**A**) Correlation analysis of DEGs between infected A549 cells. The expression levels (L2FC) of DEGs in each sample is depicted by the distribution in the diagonal. Scatter plots of the gene expression between two samples are depicted on the bottom of the diagonal. A fitted red line is indicated in red. Correlation coefficients are indicated on the top of the diagonal. P-values are indicated by asterisks. (*) p-value < 0.05. (***) p-value

